# Trophic guild and forest type explain phyllostomid bat abundance variation from human habitat disturbance

**DOI:** 10.1101/2020.02.13.946889

**Authors:** Jorge D. Carballo-Morales, Romeo A. Saldaña-Vázquez, Federico Villalobos

## Abstract

The loss of tropical forest cover caused by land-use change is causing a reduction in functional groups, such as trophic guilds. Phyllostomid bats (family Phyllostomidae) are essential in the Neotropics since they occupy up to six trophic guilds, and are pollinators, seed dispersers, and regulators of vertebrate and invertebrate populations. In this study, a series of meta-analyses were performed in order to analyse their response to habitat disturbance. Data were obtained through a comprehensive literature review whereby we measured the abundance of phyllostomid bats in disturbed habitats and conserved forests. We found that the abundance of phytophagous bats depends on the type of habitat disturbance and the type of forest where it occurs. On the other hand, animal-feeding bats reduce their abundance in any disturbed habitat regardless of disturbance type and forest type. No phylogenetic signal was found in the response of bats to habitat disturbance, nor was the response found to be dependent on the type of crop, the age of the secondary forest, or the distance to a conserved forest. These results demonstrate that feeding and the type of forest where the disturbance occurs are important aspects to understand the reduction of animal populations in the face of habitat destruction processes. This has implications on the conservation of species and their function in ecosystems.

**Highlights:**

⍰ No phylogenetic signal was found in the abundance of bats in disturbed habitats.
⍰ The abundance of bats in disturbed habitats depends on the trophic guild.
⍰ Forest altitude influences the abundance of bats in disturbed habitats.
⍰ Animal-feeding bats reduced their abundance in disturbed habitats regardless of disturbance type and forest type.

## 1. Introduction

Tropical ecosystems are losing their forest cover due to changes in land use resulting from the development of different human activities (Venter et al. 2016; Potapov et al. 2017). Urbanization, logging, agriculture, and livestock are the main land uses responsible for the current deforestation (Potapov et al. 2017). In many cases, these can cause a reduction in the abundance of species (Dirzo et al. 2014; Newbold et al. 2015) and the subsequent loss of ecosystem functions due the reduction in functional groups. For example, a reduction in trophic guilds that fulfil specific functions, such as pollination, seed dispersal, and control of populations within ecosystems, results in an imbalance of ecosystem functionality (Díaz et al. 2013; Dirzo et al. 2014; Newbold et al. 2020).

In the Neotropics, leaf-nosed bats (Phyllostomidae) are essential in ecosystem functionality. They are pollinators and seed dispersers of more than 700 plants species, and they are also predators of vertebrates and invertebrates, thus acting as regulators of their populations (Muscarella & Fleming 2007; Kalka et al. 2008; Kunz et al. 2011). The family Phyllostomidae consists of 11 subfamilies, 60 genera, and 212 species (Cirranello et al. 2016) distributed throughout the tropics and subtropics of America, including the Antilles (Villalobos & Arita 2010). Species diversity patterns of phyllostomids are considered one of the greatest adaptive radiations among vertebrate families due to the wide ecological and morphological variation they exhibit (Freeman 2000; Dumont et al. 2014). This morphological variation is reflected in six trophic guilds: nectarivores, frugivores, insectivores, carnivores, sanguivores, and omnivores (Freeman 2000; Rex et al. 2010; Rojas et al. 2011). In addition, phyllostomids have been considered good bioindicators in Neotropical forests, since their species richness and relative abundance is reduced by human habitat disturbance (Fenton et al. 1992; Medellín et al. 2000; Jones et al. 2009).

Previous studies suggest that the reduction in the populations of phyllostomid species depends on the trophic guild to which they belong because not all tropics guilds are affected in the same way by habitat disturbance (Medellín et al. 2000; Klingbeil & Willig 2009; Willig et al. 2019). In general, frugivorous and nectarivorous bats are considered tolerant to habitat disturbance, whereas insectivorous and carnivorous bats are considered sensitive. However, when analysing the response of bats from different trophic guilds in different types of habitat disturbance, contradictory results arise. This contradictory pattern has been observed in nectarivorous (Ochoa 2000; Williams-Guillén & Perfecto 2010; Durán & Pérez 2015), insectivorous (Murillo-García & Bedoya-Durán 2014), and sanguivorous phyllostomid bats (Quinto-Mosquera et al. 2013; Gonçalves et al. 2017). The latter, traditionally considered as tolerant to human disturbance activities, such as livestock, appear to be sensitive (Quinto-Mosquera et al. 2013; Gonçalves et al. 2017).

Several studies have been conducted in order to explore this controversy. It has been found that omnivorous and frugivorous bats are tolerant to livestock grassland, whereas carnivorous, insectivorous, nectarivorous, and sanguivorous bats become sensitive to this type of disturbance (Gonçalves et al. 2017). Another study that considered different types of habitat disturbance showed that carnivorous and insectivorous bats are sensitive to habitat disturbance, frugivores and nectarivores tolerate agroforestry crops, and all trophic guilds, including omnivores and sanguivores, are sensitive to monocultures and grasslands (García-Morales et al. 2013). This is consistent with the results of a recent study that found that the functional and taxonomic diversity of Neotropical bats decreases in habitats less similar to conserved forests (i.e. high contrast), such as crops, grasslands, and early-stage secondary forests (Farneda et al. 2019). However, it is not clear how bats belonging to different trophic guilds would respond to urbanization, logging, and tourism. In addition, considering the wide geographic distribution of phyllostomids, it is still unknown if the type of forest where the disturbance occurs could affect their relative abundance. Moreover, there are factors that have not been considered, such as the distance between conserved forests and disturbed habitats, and the phylogenetic signal in the response to disturbance. Previous studies have shown that the abundance of frugivorous bats decreases in wooded crops and grasslands as the distance to a patch of conserved forest increases (Galindo-González & Sosa 2003), which is similar to what has been found in birds and trees (Socolar et al. 2019). It is important to note that, because we are performing observations with bat species, it is necessary to determine the phylogenetic signal based on the phylogenetic relationship between species, since their relatedness could generate statistical non-independence (Münkemüller et al. 2012). Finally, since the last meta-analysis carried out in 2013, more than twenty studies have been conducted in the Neotropics in order to understand the response of phyllostomids to habitat disturbance. These studies can allow to make a new quantitative revision about the effect of habitat disturbance on phyllostomid abundance.

Therefore, the objective of this study was to answer, through a series of meta-analyses, the following questions:

i. Is the relative abundance of trophic guilds occupied by phyllostomids different according to the type of habitat disturbance? We expected frugivorous, nectarivorous, and omnivorous bats to be more abundant, or maintain a similar abundance, in disturbed forests, such as secondary forests, crops, mixed habitats with crops and grasslands, forests with selective logging, forests with tourism, and urban areas, compared to conserved forests. These types of human disturbances allow the presence of pioneer plants in these disturbed habitats or their surroundings, favouring phytophagous bats or those that include pioneer plants in their diet (Peters et al. 2006; Castro-Luna et al. 2007; Saldaña-Vázquez et al. 2010; Prone et al. 2012; Cisneros et al. 2015; Willig et al. 2019). We expected sanguivores to increase their abundance, or maintain a similar abundance, in secondary forests, crops, and mixed habitats with crops and grasslands, compared to conserved forests. The proximity of these habitats to human settlements and their domestic animals represents a potential source of food (Delpietro et al. 1992; Bobrowiec et al. 2015). Conversely, we expected carnivores and insectivores to be more abundant in conserved forests, compared to all other types of habitat disturbance, since deforestation promotes a reduction in prey and roosting sites (Bernard & Fenton 2003; Jones et al. 2017).
ii. Is the relative abundance of trophic guilds of phyllostomids dependent on crop intensity and secondary forest age? We expected carnivores and insectivores to be sensitive regardless of crop intensity or secondary forest age, based on our previous predictions. In the case of sanguivores, we expected their response to not be related to crop intensity or forest age, since it is believed that their response depends rather on the proximity to human settlements, as mentioned above. We expected frugivores, nectarivores, and omnivores to be tolerant to low intensity crops (e.g. agroforestry) and sensitive to high intensity crops (e.g. monocultures), since pioneer plants have been observed to be abundant in low intensity crops, and their abundance decreases as intensity increases (Williams-Guillén & Perfecto 2010). We also expected these three guilds to remain tolerant throughout the different ages of secondary forests, since pioneer plants are abundant in these habitats due to the succession process (Castro-Luna et al. 2007; de la Peña-Cuéllar et al. 2012; Farneda et al. 2018).
iii. Is the relative abundance of phyllostomid trophic guilds in disturbed habitats related to the distance to conserved forests? We expected bats, despite their ability to fly, to decrease their abundance in disturbed habitats as the distance to conserved forests increased, which is similar to patterns observed in terrestrial mammals, other bats, and birds (Galindo-González & Sosa 2003; Socolar et al. 2019; Pardo et al. 2019). We expected this because most diurnal refuges of bats are in conserved habitats (Cortés-Delgado & Sosa 2014).
iv. Is the relative abundance of phyllostomid trophic guilds related to the forest where human disturbance occurs? We expected bats to be sensitive to different types of habitat disturbance in forests with higher elevation but not in lowland tropical forests, as has been observed in another animal groups (Dalsgaard et al. 2018). This is because there is a greater availability of food in lowland tropical forests (<1000 meters), which facilitates the increase in bat populations and maintains a greater diversity of bat species compared to forests of higher elevation (Rex et al. 2008; Martins et al. 2015).

## 2. Methods

### 2.1. Literature search

We conducted an extensive review of the available literature through Google scholar. The keywords used were “phyllostomidae”, “abundancia”, “alteración”, “perturbación”, “murciélagos”, “abundance”, “perturbation”, and “bats”. We did not include words in Portuguese because most studies published in this language regularly include a title, abstract, and keywords in English. We did not limit the search by year of publication. We selected studies that sampled phyllostomids in disturbed habitats and at least one conserved forest. All the studies considered in the analysis included: number of captures of each bat species per site, distance between sampled sites, and a description of the type of habitat disturbance. When studies did not report the distance among sampled sites, we extracted it from the study site map (when present) with ImageJ 1.52a (Schneider et al. 2012).

### 2.2. Database

We obtained a total of 22 studies presenting useful data (Supporting Information). The 22 studies summarized 763 bat species observations (i.e. number of cases, k) conducted in six countries (Fig. 1). The observations comprised 107 phyllostomid bat species belonging to 42 genera and 11 trophic guilds, based on (Rojas et al. 2011), and seven types of disturbance (Table 1) and five forest types, according to the ecological zones defined by the FAO (2012).

**Table 1.** Definition of the seven types of habitat disturbance and the number of cases for each one.

| Disturbance type | Definition | Number of cases (*k*) |
| --- | --- | --- |
| Crops | All land used for growing a product regardless of its type and intensity. These were subclassified into: monoculture, polyculture, agroforestry, and urban crops. | 262 |
| Grassland | Land covered to a greater extent by grass in an induced manner and for livestock purposes. | 21 |
| Grassland-Crops | Mixed use of land by induced grassland for livestock and crops. | 20 |
| Logging | Forest intervened by selective logging. | 107 |
| Secondary forest | Secondary vegetation where, after the removal of the forest, there has been a process of natural succession. The number of years the process takes without any type of human intervention was taken. | 292 |
| Tourism | Forest intervened with the presence of trails and buildings for tourism purposes. | 7 |
| Urban zone | Sites with vegetation within the urban matrix, such as home gardens and urban parks. | 54 |

**Figure 1.**
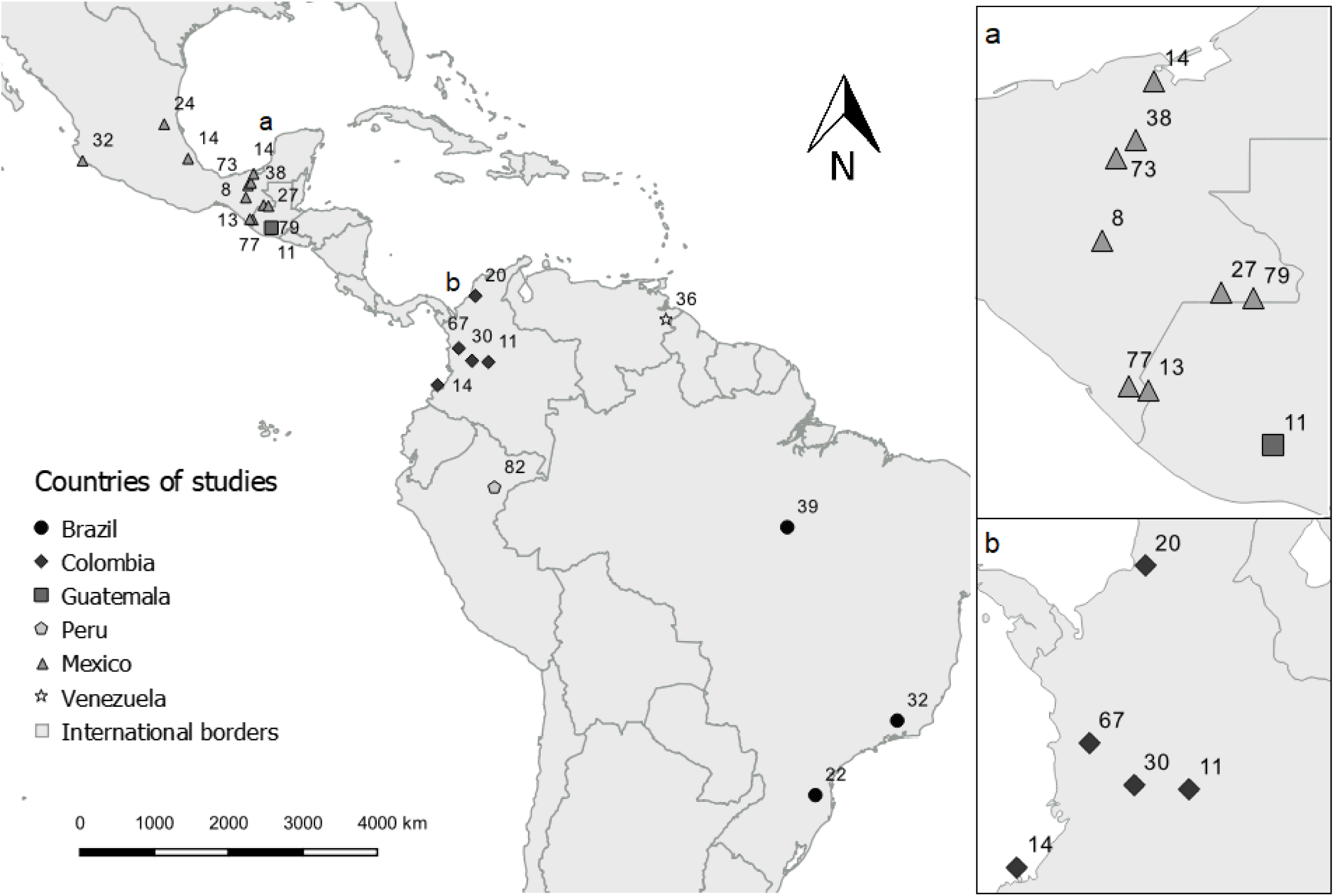
Map of the locations of the studies used for the generation of the database. Each point indicates the number of cases that contributed to the analysis. Boxes on the right show a close up of the areas marked as a and b on the main map.

### 2.3. Phylogenetic signal in phyllostomid abundance

We identified two potential sources of non-independence in our data. The first one was the possibility of phylogenetic signal in the species abundance observed in disturbed habitats (Nakagawa et al. 2017). To assess this effect, we evaluated the presence of phylogenetic signal by making a phylogenetic tree of the species present in our database. The second source was related to the author involved in the investigation (Nakagawa et al. 2017), when more than one species data item coming from the same author could be similar or biased. To deal with this effect, we included author as a random effect in our meta-analysis (see below).

For the estimation of phylogenetic signal in species abundance in disturbed habitats, we constructed a new phylogeny for the Phyllostomidae species present in our abundance dataset, since not all species with observations of abundance were present in previous Phyllostomidae phylogenies (Baker & Hoofer 2003; Datzmann et al. 2010; Rojas et al. 2011); Shi and Rabosky (2015)). Characters used were DNA sequences obtained from the Genbank database (NCBI) for Cytochrome Oxidase I (COI), Recombination activating 2 (RAG2), and Cytochrome B (CYTB) genes from 157 phyllostomid species and one mormoopid species (*Pteronotus parnellii*), which was used as an outgroup (Supporting Information). We aligned all sequences with MUSCLE using the default parameters (Edgar 2004). We performed a multilocus phylogenetic reconstruction in BEAST v1.8.4 (Suchard et al. 2018) for the Bayesian inference. We linked the trees and the molecular substitution model for all genes. We used the GTR+I+G molecular substitution model, strict clock model, and constant size model for the construction of the tree (Kingman 1982). We did three independent runs for one hundred million generations and sampled every one thousand generations. We discarded the first 2.5 million generations of each run as a burn-in. We used TRACER v1.8.2 (Rambaut et al. 2018) to estimate the effective sample size for each parameter (all resulting effective sample sizes exceeded 100) and its convergence, and to calculate the mean, upper, and lower bounds of the 95% highest posterior density interval (95% HPD). We combined the trees sampled from each independent run (10000) using LogCombiner and TreeAnnotator (Heled & Drummond 2010). The obtained grouping of species was consistent with phylogenetic trees generated in other studies (Rojas et al. 2011; Shi & Rabosky 2015).

We looked for a phylogenetic signal in the abundance of bat species by performing a randomization test in the R computational environment (R Core Team 2018). We used the obtained phylogenetic tree (Fig. 2) and its branch lengths to perform the randomization test, which evaluates the variation expected in a quantitative trait under a Brownian motion model of evolution compared with values obtained by shuffling trait data across the tips. Higher values and statistically significant values of the K index indicate a stronger phylogenetic signal (Blomberg et al. 2003). Species abundance values used in the randomization test were the mean of the proportion of bats captured in disturbed habitats for each species. To obtain these values, we performed a multivariate meta-analysis of the proportion of the abundance of species captured in disturbed habitats, taking species as a fixed factor and the author that reported the abundance value as a random factor (Viechtbauer 2010).

**Figure 2.**
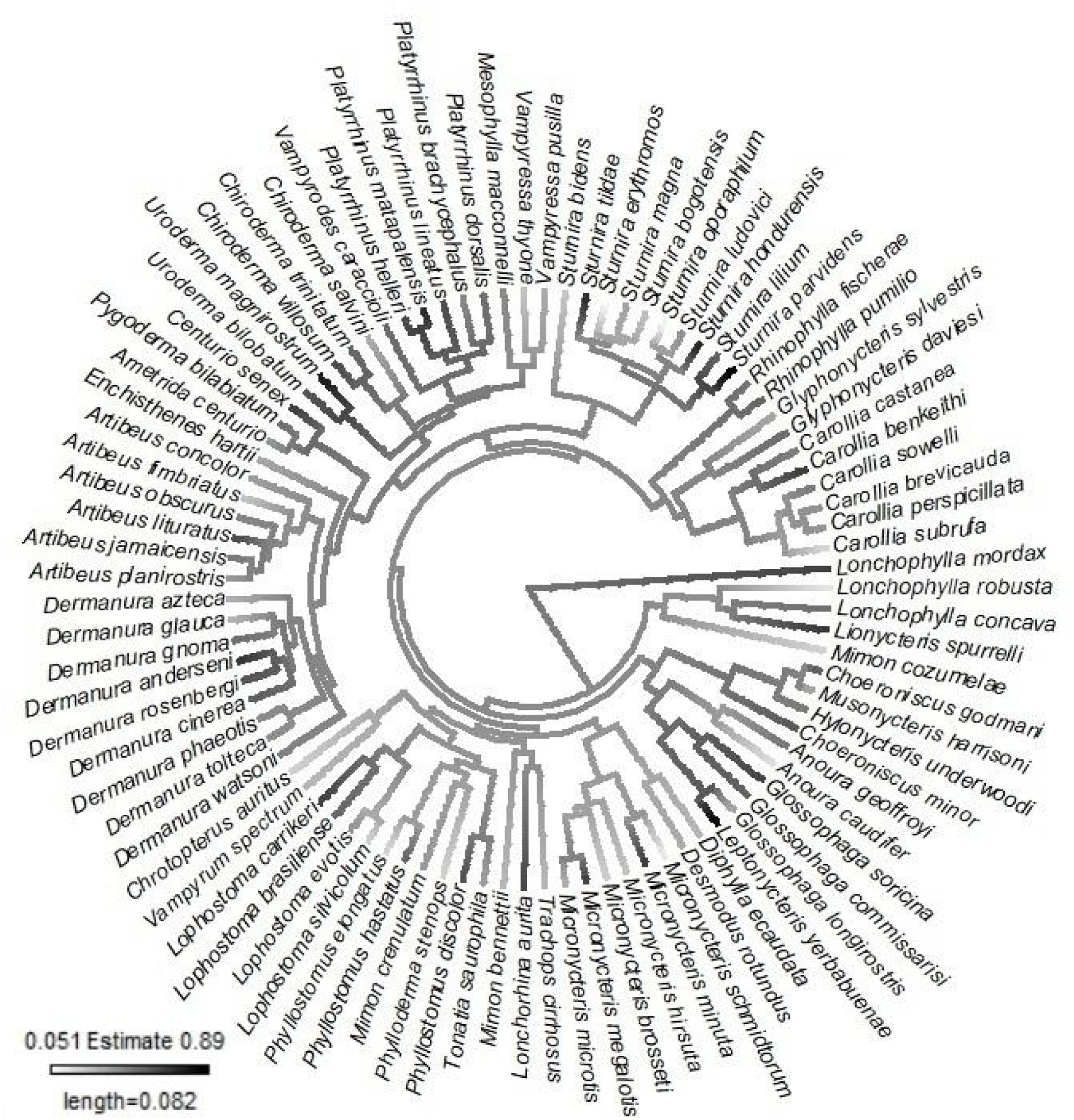
Phylogenetic reconstruction of the Phyllostomidae family using COI, RAG2, and CTBY genes. The white-black gradient in each branch is related to the proportion of the abundance of each observed species. The figure shows that there is no phylogenetic signal in the response of bats to habitat disturbance.

Given that not all phyllostomid species with abundance were present in the phylogenetic tree, we pruned the branches using the “Picante” package (Kembel et al. 2010). In this way, species in the tree corresponded to those in the database used in the meta-analysis. Of the 158 species in the tree and 107 species in the database, only 91 were used in the analysis (Supporting Information). We used the “Geiger” package (Harmon et al. 2008) to create a file that contained the species, the effect size, and the phylogenetic lengths. The phylogenetic signal was tested with the “phytools” package (Revell 2012) using 1000 randomizations. Due to the absence of phylogenetic signal in the proportion of bat abundance in the assessed species (K=0.35, P= 0.204, Fig. 2), the meta-analyses below were performed without phylogenetic correction.

### 2.4. Effect of habitat disturbance on abundance of phyllostomid trophic guilds

We performed five meta-analyses corresponding to each question in our objectives and their respective moderator variables: 1) habitat disturbance type, 2) crop type, 3) secondary forest age, 4) distance between conserved and disturbed sample sites, and 5) habitat disturbance type inside each forest type. The selected effect size was the proportion of individuals captured in the disturbed habitat from the total of individuals captured per species in both habitats (conserved and disturbed); hereafter referred to as “bat abundance”. In each meta-analysis, the author of the study was considered as a random factor (Nakagawa et al. 2017). All analyses were performed using the package “Metafor” (Viechtbauer 2010) in the R computational environment (R Core Team 2018).

We used Cochran’s Q index as a measure of heterogeneity for each analysis. Heterogeneity is important in meta-analyses because it allows to evaluate if the variation in the collected effect sizes is explained by population variation or by chance (Harrison 2011; Nakagawa et al. 2017). In addition, if heterogeneity is significant, it means that variation in effect size could be explained by moderator variables (i.e. forest type, distance, etc.). In order to examine the publication bias in our data set, we performed a regression test (Egger et al. 1997; Nakagawa et al. 2017). A significant result in the regression test indicates that effect sizes (i.e. bat abundance) are balanced. Finally, we did not perform a meta-analysis for carnivorous (C) guilds due to the low number of observations in the database (k = 2).

## 3. Results

Bat abundance in disturbed habitats was 0.46 from the total abundance observed (P<0.0001, CI=0.42-0.48, k=763), and the observations were significantly heterogeneous (Q=28237.5540, d.f.=762, P<0.0001). Therefore, the response of bats to habitat disturbance was classified as sensitive because the confidence intervals (CI) of the abundance estimate in disturbed habitats was lower than 0.5 and, thus, did not overlap with this value.

In general, we found that frugivores (F), frugivores-nectarivores (FN), insectivores-nectarivores (IN), insectivores-nectarivores-frugivores (INF), and nectarivores (N) were tolerant to habitat disturbance. On the other hand, insectivores (I), sanguivores (S), insectivores-frugivores (IF), carnivores-frugivores (CF), and insectivores-carnivores-frugivores (ICF) were sensitive to habitat disturbance (Supporting Information). In addition, for these last three trophic guilds (IF, CF and ICF), the heterogeneity of the adjusted models was not significant (Supporting Information). Therefore, we did not evaluate the effect of the moderator variables on the abundance of these guilds.

We evaluated the different moderator variables of bat abundance in disturbed habitats for bats from trophic guilds with significant heterogeneity (F, FN, IN, INF and N). We did not find differences in the abundance of these trophic guilds among different crop types (Supporting Information) despite the significant heterogeneity of the adjusted model (P <0.05, Supporting Information). Secondary forests age and distance from conserved forests were partly explained by trophic guild (□_distance_=0.1-0.5, Supporting Information); however, there was no significance in their heterogeneity (P>0.05, Supporting Information).

In the meta-analysis with habitat disturbance type as moderator, we found that only F, N, and INF responded significantly different (Table 2). We found that F were sensitive to logging, grassland, and urbanization, but tolerant to secondary forest and crops. Nectarivores were sensitive to urbanization and tolerant to secondary forest and crops. Finally, INF were sensitive to logging, grassland, and crops, but they were tolerant to secondary forest and mixed habitats such as grassland-crops.

**Table 2.** Response of frugivorous (F), nectarivorous (N), and insectivorous-frugivorous-nectarivorous (INF) bats to different types of habitat disturbance.

| Trophic guild (k) | QM<sup>a</sup> (d.f.) | Disturbance type | Estimate<sup>b</sup> | CI<sup>c</sup> | P |
| --- | --- | --- | --- | --- | --- |
| F (462) | 1019.5690 (7) | Logging<sup>d</sup> | 0.2029 | 0.0932-0.312 | 0.0003 |
|  |  | Tourism<sup>e</sup> | 0.751 | 0.634-0.869 | <0.0001 |
|  |  | Secondary forest<sup>e</sup> | 0.556 | 0.459-0.653 | <0.0001 |
|  |  | Crops<sup>e</sup> | 0.558 | 0.461-0.656 | <0.0001 |
|  |  | Grassland<sup>d</sup> | 0.215 | 0.115-0.315 | <0.0001 |
|  |  | Grassland-crops | 0.221 | 0-0.662 | 0.325 |
|  |  | Urban zone<sup>d</sup> | 0.3202 | 0.219-0.421 | <0.0001 |
|  |  | Global analysis | 0.4808 | 0.454-0.507 | <0.0001 |
| N (41) | 413.7397 (6) | Logging | 0.552 | 0.418-0.687 | <0.0001 |
|  |  | Tourism | 0.823 | 0.451-1 | <0.0001 |
|  |  | Secondary forest<sup>e</sup> | 0.785 | 0.686-0.885 | <0.0001 |
|  |  | Crops<sup>e</sup> | 0.641 | 0.522-0.7605 | <0.0001 |
|  |  | Grassland-crops | 0.5 | 0.0799-0.9201 | 0.0197 |
|  |  | Urban zone<sup>d</sup> | 0.219 | 0-0.452 | 0.0651 |
|  |  | Global analysis | 0.566 | 0.478-0.653 | <0.0001 |
| INF (26) | 504.4064 (6) | Logging<sup>d</sup> | 0.226 | 0.0683-0.383 | 0.005 |
|  |  | Secondary forest<sup>e</sup> | 0.813 | 0.712-0.913 | <0.0001 |
|  |  | Crops<sup>d</sup> | 0.359 | 0.245-0.473 | <0.0001 |
|  |  | Grassland<sup>d</sup> | 0.129 | 0-0.4707 | 0.4577 |
|  |  | Grassland-crops<sup>e</sup> | 0.962 | 0.839-1 | <0.0001 |
|  |  | Urban zone | 0.363 | 0.0013-0.725 | 0.0492 |
|  |  | Global analysis | 0.473 | 0.349-0.596 | <0.0001 |

<sup>a</sup>Cochran's Q index value for the heterogeneity test
<sup>b</sup>is the proportion of bat abundance estimated for each model.
<sup>c</sup>Confidence interval of the estimate value
<sup>d</sup>are sensitive and do not transpose their confidence intervals with <sup>e</sup>
<sup>e</sup>are tolerant and do not transpose their confidence intervals with <sup>d</sup>

Our nested analysis indicated that the response of bats depends on the forest type where the disturbance occurs only in the case of F and FN (Table 3). Frugivores were sensitive to secondary forest if it occurred in the Tropical Mountain System but tolerant when it occurred in the Tropical Rainforest. They were also sensitive to urbanization if it occurred in the Tropical Moist Forest or the Tropical Mountain System but tolerant when it occurred in the Tropical Rainforest. Frugivores-nectarivores were sensitive to secondary forest if it occurred in the Tropical Mountain System but tolerant when it occurred in the Tropical Dry Forest or Tropical Rainforest and were sensitive to crops if they occurred in the Tropical Mountain System but tolerant when they occurred in the Tropical Rainforest.

**Table 3.** Response of frugivorous (F) and frugivorous-nectarivorous (FN) bats to disturbance type in different forest types: Tropical Rainforest (TRF), Tropical Dry Forest (TDF), Tropical Moist Forest (TMF), Tropical Mountain System (TMS), and Subtropical Mountain System (STMS).

| Trophic guild (*k*) | QM<sup>a</sup> (d.f.) | Disturbance type | Forest type | Estimate<sup>b</sup> | CI<sup>c</sup> | p |
| --- | --- | --- | --- | --- | --- | --- |
| F (462) | 1213.5333 (15) | Secondary forest | TDF | 0.576 | 0.341-0.811 | <.0001 |
|  |  |  | TMF | 0.284 | 0-0.617 | 0.0943 |
|  |  |  | TMS<sup>d</sup> | 0.233 | 0-0.469 | 0.0527 |
|  |  |  | TRF<sup>e</sup> | 0.627 | 0.515-0.738 | <.0001 |
|  |  | Crops | STMS | 0.295 | 0-0.627 | 0.0818 |
|  |  |  | TMS | 0.587 | 0.349-0.826 | <.0001 |
|  |  |  | TRF | 0.588 | 0.476-0.699 | <.0001 |
|  |  | Grassland | TMS | 0.0264 | 0-0.263 | 0.827 |
|  |  |  | TRF | 0.144 | 0.028-0.2603 | 0.015 |
|  |  | Urban zone | TMF<sup>d</sup> | 0.0451 | 0-0.377 | 0.7901 |
|  |  |  | TMS<sup>d</sup> | 0.068 | 0-0.4015 | 0.6894 |
|  |  |  | TRF<sup>e</sup> | 0.6061 | 0.4098-0.8024 | <.0001 |
|  |  | Global analysis |  | 0.4808 | 0.454-0.5074 | <.0001 |
| FN (52) | 223.4907 (12) | Secondary forest | TDF<sup>e</sup> | 0.7304 | 0.485-0.975 | <.0001 |
|  |  |  | TMF | 0.75 | 0.0594-1 | 0.0333 |
|  |  |  | TMS<sup>d</sup> | 0.0385 | 0-0.388 | 0.829 |
|  |  |  | TRF<sup>e</sup> | 0.697 | 0.563-0.831 | <.0001 |
|  |  | Crop | STMS | 0.823 | 0.454-1 | <.0001 |
|  |  |  | TMS<sup>d</sup> | 0.116 | 0-0.446 | 0.489 |
|  |  |  | TRF<sup>e</sup> | 0.628 | 0.493-0.762 | <.0001 |
|  |  | Grassland | TMS | 0.0385 | 0-0.395 | 0.832 |
|  |  |  | TRF | 0.168 | 0-0.3809 | 0.121 |
|  |  | Global analysis |  | 0.524 | 0.443-0.6043 | <.0001 |

<sup>a</sup>Cochran's Q index value for the heterogeneity test
<sup>b</sup>is the proportion of bat abundance estimated for each model.
<sup>c</sup>Confidence interval of the estimate value
<sup>d</sup>are sensitive and do not transpose their confidence intervals with <sup>e</sup>
<sup>e</sup>are tolerant and do not transpose their confidence intervals with <sup>d</sup>

## 4. Discussion

Our results show that populations of phyllostomid bats are sensitive to human habitat disturbance, which contrasts with what was found in a previous meta-analysis (García-Morales et al. 2013). However, we found that this response depends on the trophic guild to which the species belong. We observed that frugivores (F), frugivores-nectarivores (FN), insectivores-nectarivores (IN), insectivores-nectarivores-frugivores (INF), and nectarivores (N) were tolerant to habitat disturbance, whereas insectivores (I), sanguivores (S), insectivores-frugivores (IF), carnivores-frugivores (CF), and insectivores-carnivores-frugivores (ICF) were sensitive. These results agree with previous observations of the effect of habitat disturbance on phyllostomid abundance (Fenton et al. 1992; Medellín et al. 2000; Ávila-Gómez et al. 2015).

Even though I and S showed significance in the heterogeneity of the adjusted models, they remained sensitive in subsequent analyses with moderator variables. The response to habitat disturbance by sensitive guilds could be explained by initial deforestation causing the loss of potential refuges. For example, S use holes in living trees as well as caves or cracks in rocks (Voss et al. 2016; Gonçalves et al. 2017), while I and C use holes in living trees or arboreal termite nests (Bernard & Fenton 2003; Kalko et al. 2006; Jones et al. 2017). In addition, I and C present fidelity and permanence towards this resource. Moreover, the prey of these species are more abundant in conserved forests (Kalko et al. 1999; de la Peña-Cuéllar et al. 2012).

### 4.1 Habitat disturbance type

Different types of habitat disturbance promote different changes in food and other resources used by phyllostomids and, thus, not all bats can tolerate disturbance in a similar way. We found F and N to have the highest variation in abundance response in relation to disturbance type. This pattern was previously observed by García-Morales et al. (2013).

We also found that F and N were tolerant to secondary forests and crops, which is consistent with previous studies (Fenton et al. 1992; Klingbeil & Willig 2009). This response may be due to the abundance of chiropterochorous or chiropterophilous species among pioneer plants in these habitats, thus favouring F and N (Castro-Luna et al. 2007; Muscarella & Fleming 2007; Castro-Luna & Galindo-González 2012). Frugivores were also tolerant to forests with tourism; however, our results are not conclusive because we only included one study with this type of habitat disturbance. This study was conducted near a conserved forest and a secondary forest, which explains the high abundance of this trophic guild (Murillo-García & Bedoya-Durán 2014). On the other hand, we expected F and N to be tolerant to urban zones and forests with selective logging, as has been observed in previous studies (Peters et al. 2006; Ferreira et al. 2010; Prone et al. 2012). However, we found the opposite, which could be explained by our analysis being performed at the trophic guild level, whereas studies that found these guilds to be tolerant were performed at the genus level (Saldaña-Vázquez et al. 2010; Saldaña-Vázquez & Schondube 2016). Finally, we found F to be sensitive to livestock grasslands, which is consistent with our expectations and the study by García-Morales et al. (2013).

### 4.2. Crop type and secondary forest age

We found that crop type and secondary forest age did not affect the abundance of phyllostomid trophic guilds. This could be explained by our study including a high number of studies on agroforestry crops versus only one on monocultures. Similarly, in the case of secondary forest age, most studies evaluated secondary forests of 15 years or less, whereas very few looked at secondary forests of 50 years or more. The results may change with data with the same number of cases per crop type according to intensity and secondary forests of different ages.

### 4.3. Distance to conserved forests

We found that the distance to conserved forests does not influence the abundance of phyllostomid trophic guilds in disturbed habitats. Other studies show the opposite effect in non-flying mammals, bats, birds, and trees (Galindo-González & Sosa 2003; Cleary et al. 2016; Socolar et al. 2019; Pardo et al. 2019). The lack of a significant effect of distance to conserved habitats on bat abundance in disturbed habitats may be related to the variation in habitat disturbance type in our study and the ability of phyllostomids to fly large distances either to migrate or forage (Arnone et al. 2016; Esbérard et al. 2017; Medellin et al. 2018). In order to determine the effect of these two variables on bat abundance in disturbed habitats, studies that evaluate the effects of habitat disturbance on bat abundance comparing disturbance type and phyllsotomid vagility are necessary.

### 4.4. Forest type

Our results show that the abundance of F and FN can vary depending on the type of forest where the habitat disturbance occurs. Both trophic guilds decreased their abundance, from tolerant to sensitive, in secondary forests of tropical mountain systems (forests with altitudes higher than 1000 meters). The same happened with FN, which became sensitive in crops located in tropical mountain systems. On the other hand, F became tolerant in urban zones when they occurred in tropical rainforests (forests with an altitude lower than 1000 meters).

The abundance and richness of phyllostomids could change according to altitude, since higher species richness and abundance has been reported in lowland forests (Sampaio et al. 2003; Rex et al. 2008), whereas abundance has been observed to decrease in habitats with altitudes above 1000 meters (McCain 2007; Martins et al. 2015; de Carvalho et al. 2019). Therefore, habitat disturbance could have a major impact on phyllostomid populations in highlands compared to lowlands. In addition, the diversity of plants is high in lowlands but decreases after 1000 meters (Gentry 1988), which explains the sensitivity of phytophagous bats to habitat disturbance in highland forests. Other studies have shown the importance of protecting high altitude tropical forests because human activities put ecosystem services at risk, for example, the protection and purification of freshwater (Martínez et al. 2009; Armenteras et al. 2011). Our results support this idea and highlight the importance of the conservation of highland tropical forests.

## 5. Conclusions

Although some bats are tolerant to habitat disturbance, because these habitats can provide food and refuges, our results do not suggest that they replace the resources provided by a conserved forest. Bats move through the matrix using both anthropic environments and conserved forests overnight (Ripperger et al. 2015). Therefore, conserved forests will always be essential to maintain phyllostomid bat populations of different trophic guilds. We conclude that the response of phyllostomid guilds to human habitat disturbance is complex and does not depend on phylogenetic signal. Sensitivity occurs regardless of disturbance type or forest type. It is advisable to enrich the anthropogenic matrix with forest cover and chiropterochorous or chiropterophilous species to promote colonization by bats and other animals in order to facilitate their functions and ecosystem services (Kunz et al. 2011; Castro-Luna & Galindo-González 2012).

The literature used in this meta-analysis was limited because some published studies do not describe the type of habitat disturbance. Also, some studies do not use a control site (conserved forest) and they do not show a map or coordinates of the sampled sites. We highlight the absence of studies in Central America and the Antilles, which is partly due to the limitations already mentioned. For future research, it is necessary to have more information about phyllostomid species variables, such as body mass, flight strategies, foraging range, and type of refuge used. These can be decisive in understanding the complex response of phyllostomid guilds to habitat disturbance.

## Appendix A. Supplementary data

The supplementary data to this article are available online.

